# On the Mental Status of Stargazer Mice

**DOI:** 10.1101/2022.02.21.481350

**Authors:** Catharina Schirmer, Mark A Abboud, Samuel C Lee, John S Bass, Arindam G Mazumder, Jessica L Kamen, Vaishnav Krishnan

**Affiliations:** Department of Neurology, Baylor College of Medicine, Houston, TX USA

## Abstract

In many childhood-onset genetic epilepsies, seizures are accompanied by neurobehavioral impairments and motor disability. In the Stargazer mutant mouse, genetic disruptions of *Cacng2* result in absence-like spike-wave seizures, cerebellar gait ataxia and vestibular dysfunction. Here, we combine videotracking and instrumented home-cage monitoring to resolve the neurobehavioral facets of the murine Stargazer syndrome. We find that despite their gait ataxia, stargazer mutants display horizontal hyperactivity and variable rates of repetitive circling behavior. While feeding rhythms, circadian or ultradian oscillations in activity are unchanged, mutants exhibit fragmented sleep bouts, atypical licking dynamics and lowered sucrose preference. Mutants also display an attenuated response to visual and auditory home-cage perturbations, together with profound reductions in voluntary wheel-running. Our results reveal that the seizures and ataxia of Stargazer mutants occur in the context of a more pervasive behavioral syndrome with elements of encephalopathy, repetitive behavior and anhedonia. These findings expand our understanding of the function of *Cacng2/CACNG2*, variants in which have been identified in patients with bipolar disorder and schizophrenia.

## Introduction

In patients with developmental and/or epileptic encephalopathies (DEEs/EEs)^1^, seizures and static intellectual disability are often accompanied by motor impairments (quadriparesis, ataxia), disturbances in sleep/arousal, sensory integration, feeding and emotional lability.^2, 3^ Behavioral comorbidities in particular amplify the risk of psychiatric side effects to antiseizure medications^4^, and compound polypharmacy through concurrent prescriptions for neuroleptics, psychostimulants and/or sedatives. With advances in mouse genome engineering, patient-informed genetic DEE mouse models now play a dominant role in devising precision genetic treatments, informed by the latest advances in transcriptomics, neuroimaging and neurophysiology^5-7^. In contrast, preclinical endpoints to ascertain neurobehavioral impairment have not experienced similar technical or conceptual innovations. Today’s most exciting mouse models often remain subject to a battery of maze- or arena-based assays (e.g., open field test, elevated plus maze, etc.)^8-10^. While enormously popular, these tests produce snapshots of behavior in singular readouts that may be amplified/attenuated by arena novelty, and which may be contaminated by variations in motor drive (e.g., hyperactivity) or function (e.g., ataxia).^11^ A growing movement seeks to adjunct these assays with naturalistic, continuous and unbiased assessments of spontaneous self-motivated behavior that can be analyzed across multiple time scales.^12-15^ In this new paradigm, prolonged experimenter-free home-cage recordings minimize observer effects^16, 17^, and automated data collection enables computational phenotypes that can be appraised and compared without anthropomorphization.^13, 14, 18^

Here, we apply one such home-cage monitoring platform^12, 19-21^ to visualize the extent and severity of behavioral impairment in Stargazer mutant mice^22^. These mutants arose from a naturally occurring disruptive transposon insertion in *Cacng2* encoding STARGAZIN, a transmembrane protein necessary for AMPA-subtype glutamate receptor expression. Characteristic 6-9Hz spike/wave seizures in mutants beginning in adolescence^22-25^ have been linked to reduced AMPAR expression in feed-forward inhibitory thalamocortical neurons.^26, 27^ Mutants also display gait ataxia, severely impaired rotarod performance and are unable to swim^28^: motor and vestibular phenotypes have been linked to diminished AMPAR expression in cerebellar Purkinje cells^27^ and/or vacuolar degeneration of the vestibular epithelium.^28^ Using prolonged experimenter-free recordings and provocative maneuvers applied *within* the home-cage, we innovate a set of fresh endpoints to define and resolve a novel syndrome of neurobehavioral impairment in Stargazer mutants.

## Methods

Protocols were approved by the Baylor College of Medicine Institutional Animal Care and Use Committee and conducted in accordance with USPHS Policy on Humane Care and Use of Laboratory Animals. Heterozygous mutant mice were bred to obtain wildtype (WT) and homozygous mutant mice. PCR-based genotyping was performed on tail DNA at ~p16^23, 24, 29^, and mice were weaned into gender-matched cages at p21. Between ~p21-28, all weaned cages were provided with Bio-Serv Nutragel as to supplement nutrition to mutants in the setting of ataxia and low body weight. At ~p50, mice were transferred to Noldus Phenotyper ^®^ home-cages (30×30×47cm) within a designated satellite study area^19, 20^. Each cage contains (i) two lickometered water sources (0.8% sucrose-drinking water Vs drinking water), (ii) an infrared(IR)-lucent shelter, an aerial IR camera and IR bulb arrays, (iii) a beam-break device to measure entries into a food hopper, (iv) a detachable running wheel, and (v) a 2300Hz pure tone generator and an LED house light. Satellite temperature (20-26C), humidity (40-70%) and light cycle settings (ON between 0500-1700) matched vivarium conditions. White noise was played continuously, and satellite access was restricted to gowned, gloved, masked and capped personnel to minimize olfactory variations.

Live videotracking (Noldus Ethovision XT14) sampled object x-y location (“centerpoint”) at 15Hz, providing time series data for arena position (heat maps, shelter time), horizontal displacement (velocity) and relative angular velocity (*positively* signed for counterclockwise turn angles). “Sleep” epochs were defined as contiguous periods of immobility lasting ≥40s, previously validated to provide >90% agreement with neurophysiologically-determined sleep.^30-33^ WT (n = 24, 9 female) and mutants (n = 26, 16 female) were studied in cohorts of 8-15 mice. A modular design^19^ was applied beginning with a 2h-long initial habituation study (“Intro”) and two consecutive 23h long baseline recordings each beginning at 1400. Then, we applied visual and auditory stimulation (“light spot^19, 20, 34^ and beep^19^”), followed by a third prolonged recording in the presence of a running wheel. We concluded with two daytime provocations to interrogate their response to transient removal of their shelter (“shelter removal”) and a novel cage stress (“cage swap”).

For simultaneous electroencephalography (EEG), six ~7-week old mutants were implanted with EMKA easyTEL S-ETA devices under sterile precautions and isoflurane anesthesia. Biopotential leads (2) were affixed subdurally in right frontal and left posterior parietal regions using dental cement, with wires tunneled to a transponder positioned in the subject’s left flank. Wireless EEG was acquired at 1000Hz sampling rate with IOX2 software (EMKA Technologies) via easyTEL receiver plates placed underneath home-cages, and EEG signals were inspected with LabChart reader using a bandpass filter (1-30Hz). Seizures were defined as bursts of 7-9Hz spike/wave discharges with an amplitude at least 2x baseline voltage.^35^ Data were graphed and analyzed with Prism Graphpad 9, always depicting mean ± standard error of the mean. Lomb-scargle periodograms (Matlab) were applied to calculate the power and peaks of ultradian oscillations in activity. Two-tailed, unpaired student’s T tests were applied, with *, **, ***, **** depicting p<0.05, <0.01, <0.001 or <0.0001 respectively.

## Results

On admission to home-cages at ~p50, mutants were comparatively underweight. Their initial habituation response was marked with hyperactivity and reduced shelter engagement (Figure 1A). A wobbling titubating gait was plainly evident. Some mutants engaged in repetitive circling behavior (Supplemental Movie 1), producing circular tracks of varied diameter that were not necessarily concentric. This resulted in an overall increase in *positive* total net angular displacement (NAD), reflecting a preference for counterclockwise circling. Absolute NADs remained significantly higher in mutants even after normalizing by total distances, with mutants accumulating a net rotation of ~± *31deg* for every centimeter of horizontal displacement (vs *4.9 deg/cm* in WT mice, Figure 1B). While both groups thoroughly explored non-shelter regions (Figure 1C), mutants also displayed a relative avoidance of waterspouts and their food hoppers (Figure 1D, E). Together, these results define the initial mutant response to an enriched novel cage, marked by hyperactivity, repetitive circling behavior and novel object avoidance.

**Figure 1:**
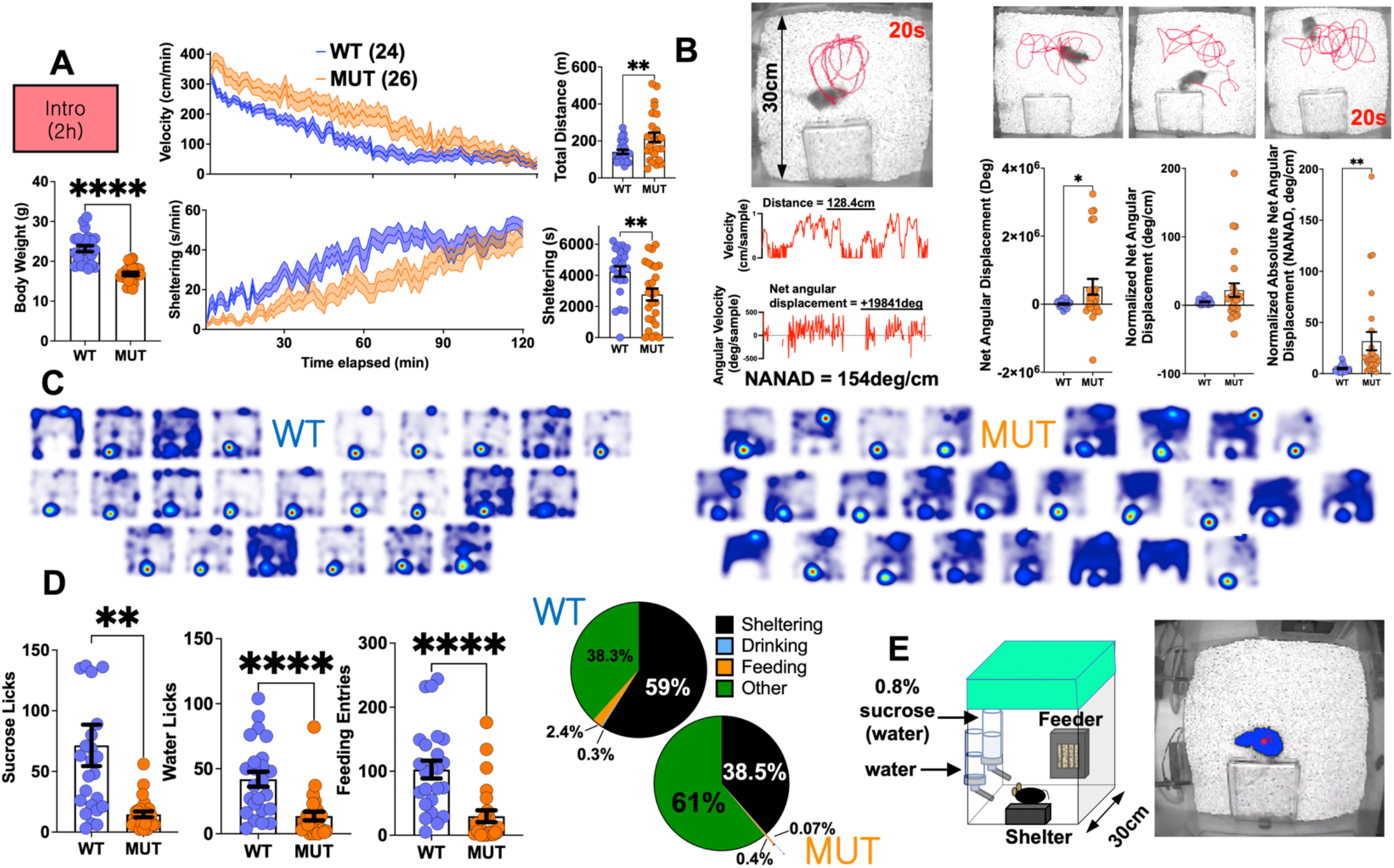
Habituation to the home-cage. A, In a 2h-long introductory trial, mutants displayed hyperactivity and reduced shelter engagement. B, Many mutants displayed repetitive circling behavior. LEFT: An illustration from a circling mouse depicting a 20s-long track, where a horizontal displacement of ~128cm is associated with a ~ +19841 deg angular displacement (signed +ve for counterclockwise turns), resulting in a [distance] <u>N</u>ormalized <u>A</u>bsolute <u>N</u>et <u>A</u>ngular <u>D</u>isplacement (NANAD) of ~154 deg/cm. RIGHT: Averaged across the entire trial, mutants displayed significantly higher NANAD values. C, Heat maps of position probability during the 2h trial. D, Mutants accumulated significantly fewer licks and feeding entries, alternatively visualized as time budgets (right). E, Schematic of home-cage design and representative aerial view with centerpoint (red) and body contour (blue). Mean ± s.e.m shown.

During subsequent baseline recordings, mutants remained hyperactive predominantly during the nocturnal phase, with a similar spectral distribution of ultradian oscillations in locomotor behavior (Figure 2A, B). Velocity raster plots (cm/min) revealed generally similar “bursty”^15^ patterns of rest and activity in both groups, with greater synchrony at dark-light transitions. Measures of circling behavior poorly correlated with total daily distances (Figure 2C). To more granularly examine changes in arousal, we applied an established and validated noninvasive method which estimates “sleep” as periods of sustained immobility lasting ≥ 40s.^19, 20, 30-33^ Under these criteria, mutants displayed shorter and more frequent bouts of “sleep”, without an overall change in total “sleep”. These differences primarily affected the ~35% of all “sleep” bouts longer than ~230s, while the timing of “sleep” bouts throughout the day was unchanged (Figure 2D). In a separate group of mutant mice monitored wirelessly with EEG, we observed that spike/wave seizures were typically brief (2-6s) and seen primarily during (without prolonging) bouts of “sleep” (Figure 2E).

**Figure 2:**
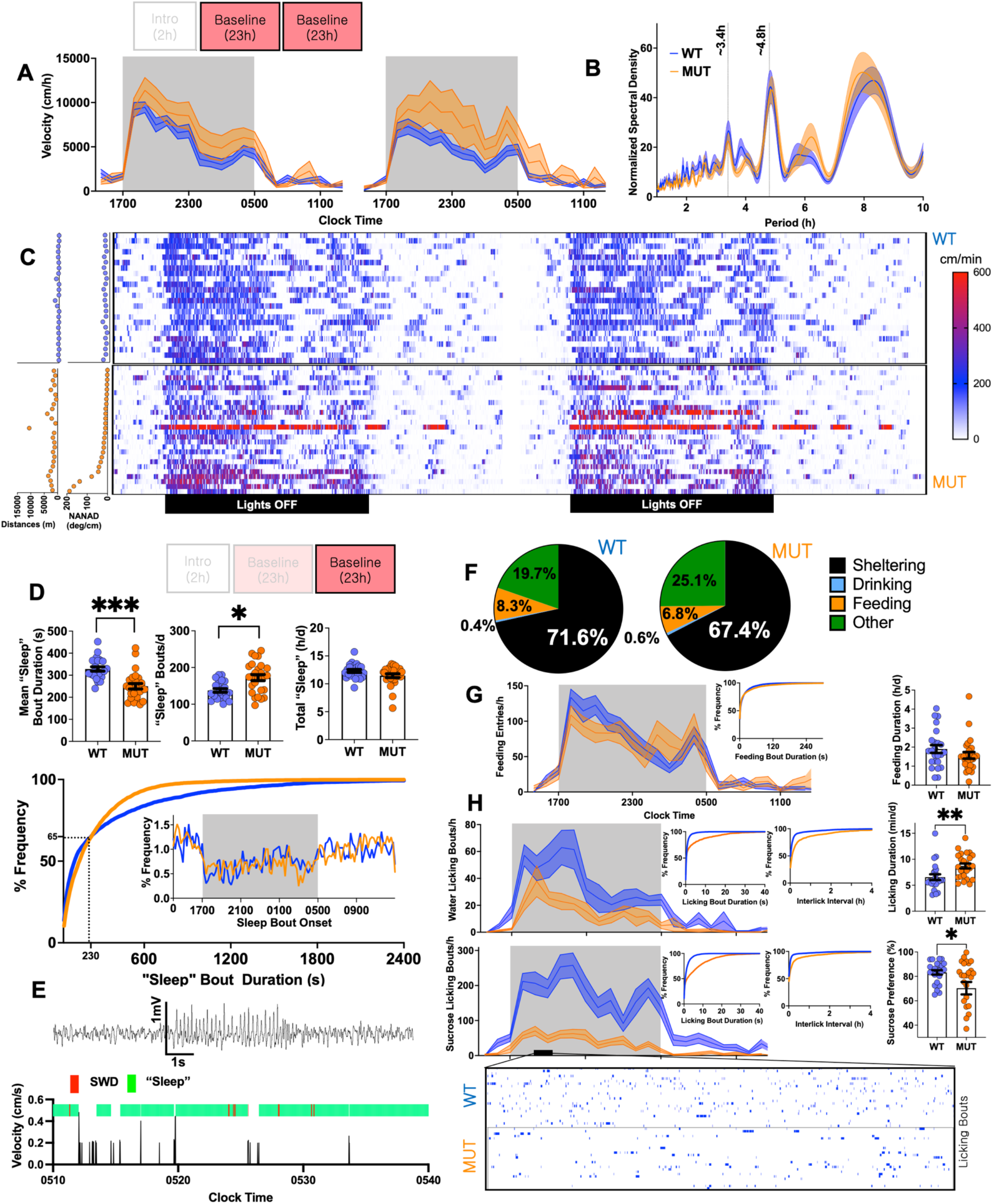
A representative day. A, During 2 consecutive 1400-1300 baseline recordings, mutants navigated greater horizontal distances. B, Lomb-scargle periodogram of ultradian oscillations in locomotor activity depicting peaks at harmonic frequencies of the circadian oscillator. C, Raster plot of individual velocities over two baseline recordings. Hyperactivity was not directly related to high measures of circling (NANAD). D, Mutants displayed fewer but longer “sleep” bouts without affecting total daily sleep. BOTTOM: Frequency distribution histogram comparing “sleep” bout durations and bout onset (inset). E, Wireless EEG conducted within home-cages revealing that spike/wave discharges occurring within “sleep” bouts. F, Daily time budgets, revealing lower shelter times and increased “other” behavior. G, Mutants exhibit total feeding times, feeding entries and bout durations that are similar to WTs. H. Mutants accomplish fewer but longer licking bouts of sucrose or water. Separately, mutants also displayed an overall reduction in sucrose preference. Mean ± s.e.m shown.

Next, we tallied total durations of licking, feeding and sheltering during the second baseline recording to visualize averaged daily time budgets^36^. Mutants displayed a trend towards lower sheltering times (p=0.1), which could not be explained by increased feeding (Figure 2F, G). Lickometry data revealed major abnormalities in the structure of fluid intake, with mutants accomplishing *greater* overall licking durations through a combination of *fewer* but *longer* licking bouts. Simultaneously, while both groups preferred sucrose water, mutants displayed a significantly lowered sucrose preference (Figure 2H). Collectively, these baseline data highlight a set of diffuse alterations in the spatiotemporal structure of spontaneous behavior in mutant mice.

We began our provocative maneuvers with an hour-long “light spot test”^19, 20, 34^, which poses a conflict between nocturnal foraging and light aversion within their home-cage and without human presence. WT mice responded acutely with *increased* activity followed by sustained immobility within shelters. In comparison, a relatively blunted locomotor suppression response was seen in mutants, with some minimal persistent exploration of the open illuminated cage (Figure 3A). Similarly, in response to a 60s-long pure tone, mutants exhibited an attenuated startle and sheltering response compared with WT mice (Figure 3B). On the following day, when provided access with a running wheel, wheel-running behavior was almost entirely absent in mutants, with only 2/26 mutants successfully engaging the wheel. Compared with baseline recordings, wheel access suppressed feeding in WT mice while this effect was absent in mutants (Figure 3C).

**Figure 3:**
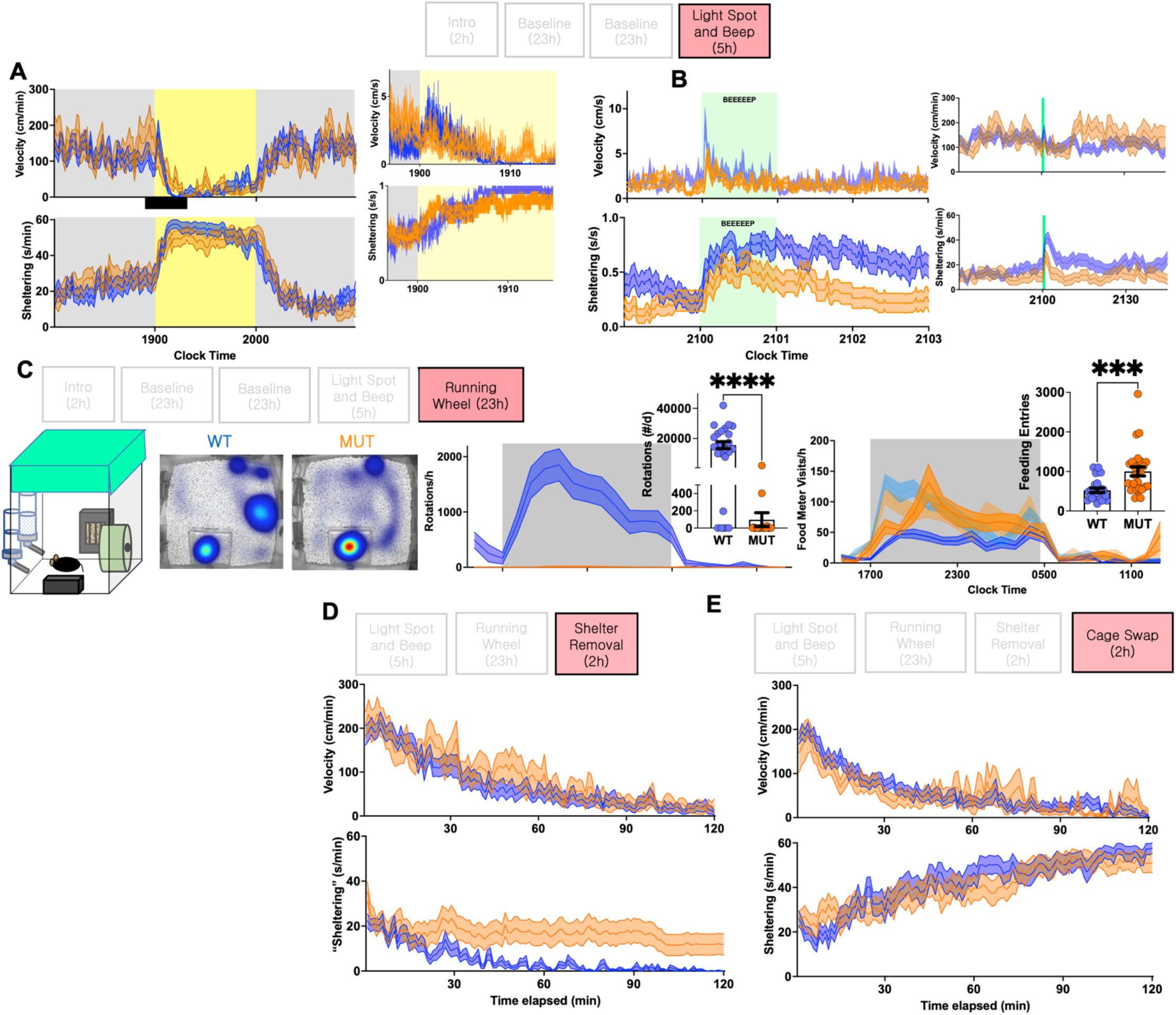
Provocative maneuvers. A, Responses of WT and mutants to the “light spot test”. B, Mutants display an attenuated early locomotor startle and sheltering response to a 60s long home-cage beep stimulus. C, Mutants display a significant reduction in wheel-running behavior. Unlike mutants, WT-mice suppress feeding entries in the presence of the running wheel (baseline feeding entries are shown as light blue, light orange). D, Locomotor and sheltering response to shelter removal, and E, cage swap protocol. Mean ± s.e.m shown.

Finally, we applied two additional brief daytime perturbations. Since mice spend the bulk of their light period/daytime within shelters^19, 20^, we examined their response to transient shelter removal. While WT mice settled in cage corners, many mutants rested in the cage location previously occupied by the shelter (Figure 3D). In a second daytime maneuver, mice were manually repositioned into a cage previously inhabited by a gender-matched mouse, providing a geometrically similar field with novel olfactory cues. Unlike their initial habituation response (Figure 1A), activity patterns and gradual shelter entry were similar (Figure 3E).

## Discussion

Advances in wearable technologies are poised to radically transform the diagnosis and management of mental health symptoms by supplementing subjective and qualitative clinic-based assessments of mental status with more continuous, *in situ* and quantitative device-based measurements.^37^ The home-cage approach provides a preclinical platform to align basic neuroscience with the digitization of neuropsychiatric symptom . This strategy recognizes that behavioral symptom constellations in mice may not (and need not) perfectly parallel human DSM-defined symptom clusters (e.g., autism spectrum disorder). By incorporating a wide array of behavioral endpoints, unique models of pervasive neurodevelopment can reveal unique constellations of behavioral impairment. For example, we have previously shown that mice with *Scn1a* mutations (modeling Dravet syndrome) display nocturnal hypoactivity, hypersomnia and hypodipsia,^20^ while mice exposed prenatally to valproic acid exhibit hypophagia and increased wheel-running.^19^

Here, we report that the seizures and ataxia of Stargazer mutants are accompanied by a behavioral syndrome reflective of broad and bilateral cerebral network dysfunction, principally featuring sleep fragmentation, anhedonia, hyperactivity and altered sensorium. Through lickometry, we find that mutants also display polydipsia, with *more* prolonged but *less* frequent licking bouts. This phenotype may reflect a synergy between cerebellar dysfunction (*licking ataxia*) and an underlying limbic disturbance in consumptive behavior (*psychogenic polydipsia*^38^). About a third of mutant mice exhibit repetitive circling behavior^28^, a phenomenon observed in rodents with lateralized asymmetries of nigrostriatal dopamine signaling, vestibular disease^39^, and in several genetic mouse models of autism spectrum disorder.^40^ Despite their hyperactivity, the vast majority of mutants do not engage in wheel-running. This may relate to cerebellar ataxia, vestibulopathy and/or anhedonia.

It remains to be determined whether some or all aspects of this syndrome may remit with early long term antiseizure treatments, which can themselves produce marked changes in home-cage behavior in wild type mice.^19^ In one previous study, ethosuximide treatment successfully suppressed spike/wave seizures in Stargazer mutants without impacting underlying aberrant patterns of EEG phase-amplitude coupling^23^, providing at least one correlate of a static network derangement that is unaffected by temporary seizure suppression. *CACNG2* genetic variants have been identified in small studies of patients with schizophrenia and bipolar disorder (without epilepsy or ataxia)^41-44^, supporting a broad role for *CACNG2* in shaping cognitive and emotional behaviors even in the absence of seizures.

In conclusion, our study provides an unbiased behavioral characterization of the Stargazer mutant mouse using a novel approach that can be applied broadly to study mice that model a triad of seizures, neurobehavioral impairment and motor abnormalities. With similar analyses, future studies may discern how these behavioral endophenotypes may be affected in mutants with graded variations in *Cacng2* expression (*Stargazer-*3J mice, *Waggler*) or in mutants with etiologically distinct syndromes of ataxia and absence seizures (*lethargic [Cacnb4], ducky [Cacna2d2], tottering [Cacna1a]*).^45^

## Acknowledgements

VK acknowledges research funding from the NINDS (1K08NS110924), an American Epilepsy Society Junior Investigator Award (2020) and seed funding from the Baylor College of Medicine Office of Research.

## Disclosure of Conflicts of Interest

VK’s laboratory has received support from SK Life Science for contract research unrelated to the scope of this publication.

## Supplemental Data

Supplemental Movie 1: Representative recordings from WT (top and bottom left) and MUT (top and bottom right) mice captured during the introduction trial. The mutant mouse on the bottom right engages in repetitive circling behavior ~35s into the recording.

## Notes

### Competing Interest Statement

The authors have declared no competing interest.

